# Transcriptome and metabolome analyses reveal novel genetic targets for L-tryptophan overproduction in *Corynebacterium glutamicum*

**DOI:** 10.1101/2025.05.07.652659

**Authors:** Yufei Dong, Rongsheng Gao, Nan Qin, Kunyu Liu, Youmeng Liu, Zhen Chen

## Abstract

*Corynebacterium glutamicum* is a promising microbial chassis for the industrial production of L-tryptophan, which has exhibited increasing demand due to its diverse applications and high market value. In previous work, we developed an L-tryptophan-overproducing *C. glutamicum* strain TR26 through multiple rounds of rational metabolic engineering. Here, comparative transcriptome and metabolome analyses were conducted between TR26 and its progenitor strain MB001 to reveal the underlying mechanisms and potential bottlenecks for L-tryptophan production in TR26. Furthermore, by systematically down- and up-regulating differentially expressed genes of interest, two novel genetic targets, *glnK* and *sugR*, were identified as being associated with L-tryptophan synthesis. Specifically, the repression of *glnK* and overexpression of *sugR* in strain TR26 increased the titer of L-tryptophan by 6.7% and 20.9%, respectively. Gene transcription profiling and intracellular metabolite analysis further suggested that the observed improvements in L-tryptophan synthesis could be attributed to optimized nitrogen transport and metabolism, efficient reallocation of cellular resources and enhanced supply of phosphoenolpyruvate (PEP). This study advances our understanding of the regulation mechanisms governing L-tryptophan synthesis in *C. glutamicum* and provides valuable insights for further optimization of industrial cell factories.

## 1. Introduction

L-tryptophan, an essential amino acid for human and animals, plays vital roles in both protein synthesis and the regulation of physiological functions. L-tryptophan has been extensively applied in pharmaceutical, nutraceutical, food and feed industries, demonstrating substantial economic value [1,2]. Over the past few decades, microbial fermentation has emerged as the most widely adopted method for L-tryptophan production, excelling other approaches due to its high efficiency, lower costs, and environmental friendliness. Industrial hosts, including *Escherichia coli* and *Corynebacterium glutamicum*, have been successfully engineered for L-tryptophan production through various strategies, such as enhancing the L-tryptophan synthesis pathway [3-7], optimizing precursor supply [8-11], implementing transport engineering [12-17], and blocking competing pathways [18,19]. Leveraging well-developed genetic manipulation tools, *E. coli* strains have been engineered to achieve L-tryptophan titers ranging from 40 to 55 g/L, with yields between 0.15 and 0.23 g/g glucose [8,15,20]. In comparison, *C. glutamicum* offers superior safety, enhanced industrial robustness, and capability to utilize various substrates for efficient amino acids production [21-28]. By integrating random mutagenesis and rational design strategies, a *C. glutamicum* strain capable of accumulating 58 g/L L-tryptophan was developed, highlighting its potential for industrial production [11]. Despite these advancements, the L-tryptophan yields achieved by both chassis organisms remain far below the theoretical maximum, which is possibly due to inadequate engineering of key metabolic nodes. Furthermore, limited understanding of L-tryptophan synthesis regulation, particularly in *C. glutamicum*, has hindered rational engineering efforts aimed at increasing product accumulation. Therefore, the identification of novel genetic targets and deeper elucidation of the L-tryptophan biosynthesis regulation mechanisms are essential for developing industrial strains with higher production capacities.

Multi-omics technologies have emerged as powerful tools for understanding the global gene expression and metabolic characteristics of industrial strains. Recent advances in next-generation sequencing techniques have substantially enriched transcriptome databases, enabling comprehensive analysis of important industrial traits such as hyper-production and stress tolerance, providing new insights into cell metabolism and regulation networks [29-35]. Additionally, novel genetic targets can be identified through transcriptome analysis, facilitating further optimization of industrial strains. For example, by analyzing the transcription profile of an L-tryptophan-producing strain, transcription factor YihL was identified to negatively regulate L-tryptophan synthesis in *E. coli* [16]. The deletion of *yihL* improved the product titer by 16.3% [16]. Similarly, transcriptional profiling was employed to decipher the mechanisms of L-phenylalanine overproduction in *E. coli*, revealing MarA as a key regulator that enhances cell tolerance to the product [36]. On the other hand, metabolome analysis complements these efforts by providing global carbon distribution profiles, which is particularly effective in identifying intermediate accumulations and bottlenecks for product accumulation [17,37,38].

In our previous work, we established the L-tryptophan overproducing *C. glutamicum* strain TR26 using rational metabolic engineering strategies [17]. In this study, we conduct comparative metabolome and transcriptome analyses of TR26 and the wild-type strain MB001. By examining the intracellular concentrations and global gene expression characteristics of these strains, we systematically elucidate the critical mechanisms underlying L-tryptophan overproduction. In addition, by down- and up-regulation of the key differentially expressed genes (DEGs), we identify novel genetic targets for L-tryptophan production in *C. glutamicum*. Further investigation into intracellular metabolites and transcriptional profiling clarifies the regulation mechanisms of these new targets. This work improves our understanding of L-tryptophan synthesis regulation in *C. glutamicum*, offering valuable insights for strain engineering towards industrial application.

## 2. Materials and methods

### 2.1. Bacterial strains and plasmids

*C. glutamicum* MB001 was a prophage-free strain derived from ATCC 13032 [39]. *C. glutamicum* TR26 was an L-tryptophan-overproducing strain derived from strain MB001 through multiple rounds of genetic engineering [17]. *E. coli* DH5α was used as the host for plasmid construction. The pD9SG plasmid [40], an *E. coli* and *C. glutamicum* shuttle vector containing dCas9 protein and sgRNA cassette sequences, was employed for gene expression repression in *C. glutamicum*. The pEC-K18mob2 plasmid [41], an *E. coli* and *C. glutamicum* shuttle vector, was used for gene overexpression in *C. glutamicum* strains.

### 2.2. Plasmids and strains construction

To construct plasmids for gene expression inhibition, the gene-specific sgRNA fragments with BsaI sticky ends were obtained via the annealing of two specific oligonucleotides, and subsequently ligated into the BsaI sites of the plasmid pD9SG using T4 ligase. The sgRNA sequences targeting each candidate gene are listed in Table S1.

To construct plasmids for gene overexpression, the functional genes with a common ribosome binding site (aaaggaggttgtc), and the rrnB terminator, were inserted into the restriction sites EcoRI/XbaI of the plasmid pEC-K18mob2 via Gibson assembly following the standard procedures. Gene expression levels were controlled by the native lac promoter on the plasmid backbone. The primers utilized for amplification of the target genes are listed in Table S2. Primers PEC_F (gtggctgttttggcggatgag) and rrnB_R (aagcttgcatgcctgcaggtcgactagagtttgtagaaacgcaaaaaggcc) were utilized to amplify the rrnB terminator seuqnece.

### 2.3. Medium and culture conditions

For shake-flask fermentations, *C. glutamicum* strains were initially cultured in 20 mL LSS1 medium in 250 mL baffled shake flasks for 14-16 h [46]. The seed cultures were subsequently inoculated into 30 mL LPG2 medium in 500 mL baffled shake flasks with 30 g/L of CaCO_3_ for pH maintenance to perform L-tryptophan fermentations [47]. The initial glucose concentration in the fermentation medium was 100 g/L. All shake-flask fermentations were conducted in duplicate at 30 °C, 200 rpm and an initial pH of 7.2.

Microtiter fermentations were conducted in Biolector^®^ I in 48-well microtiter plates. *C. glutamicum* strains were first cultured in 1 mL modified LSS1 medium with 20 g/L 3-Morpholinopropanesulfonic acid (MOPS) instead of CaCO_3_ for pH maintenance for 12-14 h. The seed cultures were then inoculated into 1 mL modified LPG2 medium, which contained 50 g/L glucose and 20 g/L MOPS without CaCO_3_ addition, and cultured for 36 h to perform L-tryptophan fermentation. All microtiter fermentations were conducted in duplicate at 30 °C, 990 rpm, and 75% relative humidity. When necessary, 25 μg/L kanamycin was added. Additionally, 1 mmol/L IPTG was included to induce the expression of the dCas9 protein for target gene repression.

### 2.4. Analysis of cell growth and extracellular metabolites

Cell growth was assessed by measuring the optical density of 600 nm (OD_600_) of the cultures. The concentrations of glucose and acetic acid were determined using high-performance liquid chromatography (HPLC) equipped with an Aminex HPX-87H Column (300×7.8 mm). The mobile phase consisted of 5 mM H_2_SO_4_, which a flow rate of 0.8 mL/min at 65 °C. The injection volume was 20 μL.

The concentrations of L-tryptophan were analyzed by HPLC with a Dikma Diamonsil AAA Column (5μm, 250×4.6 mm) using the standard phenyl isothiocyanate derivative method [48]. L-tryptophan was separated using mobile phase A (50 mM CH_3_COONa, pH 6.5) and mobile phase B (methanol: acetonitrile: water=1: 3: 1) with gradient elution. The total flow rate was 1 mL/min at 45 °C. The detective wavelength was set at 254 nm and the injection volume was 10 μL.

### 2.5. Metabolome analysis

Cells were cultured to the exponential phase and 2 mL of each sample were taken. The cell suspension was centrifuged for 1 min at 4 °C, 12000 rpm to remove the medium. The cell pellets were washed gently three times with PBS buffer (pH=7.2-7.4) and the supernatant was removed. Immediately, 1.6 mL of pre-cooled (-80 °C) 80% methanol was added to resuspend the cell pellets by vortex. The samples were then incubated at -20 °C for 30 min, followed by ultra-sonication for 2 min. Afterwards, the samples were incubated at -80 °C for 2 h for cell lysis. The suspension was subsequently centrifuged at 4 °C, 14000 rpm for 15 min to remove the protein. The supernatant was transferred to a new 2 mL microcentrifuge tube, dried by vacuum and stored at -80 °C for further operation.

The metabolites in the supernatant were analyzed by using SCIEXTriple Quad 6500+ liquid chromatography-tandem mass spectrometry (LC-MS/MS) system (AB Sciex, Singapore) equipped with IonDrive detector and Qtrap-6500 mass spectrometer. Metabolites with significantly concentration variations were identified based on the criteria: average fold change ≥1.5, t-test p ≤0.05.

### 2.6. Transcriptome analysis

Cells were cultured to the exponential and stationary phases, and 2 mL of each sample were collected. The cell suspension was centrifuged at 4 °C, 12000 rpm for 1 min to remove the medium. The cell pellets were subsequently washed three times with PBS buffer (pH=7.2-7.4) and fast-frozen in liquid nitrogen for subsequent analysis. RNA extraction and transcriptome analysis were performed by AZENTA life science. Transcriptome sequencing was conducted using the Illumina platform, with the obtained data evaluated and filtered by FastQC (v0.10.1) and Cutadapt (v1.9.1). Software bowtie2 (v2.2.6) was used to align the reads to the reference genome of *C. glutamicum* MB001 (CP005959.1). Differential expression analysis was performed using the DESeq2 Bioconductor package. Significantly differentially expressed genes were identified following the criteria: average fold change ≥2.0, t-test p ≤0.05.

## 3. Results and discussion

### 3.1. Metabolome analysis of *C. glutamicum* TR26 and MB001

*C. glutamicum* TR26 is an L-tryptophan-overproducing strain derived from the wild-type MB001 through multiple genome modifications guided by rational engineering methods. To investigate the underlying mechanisms and potential bottlenecks for L-tryptophan synthesis in TR26, comparative metabolome analysis was conducted between the TR26 and MB001, focusing on variations in the intracellular concentrations of central metabolic intermediates and amino acids. As shown in Fig. 1A, 75 metabolites in central metabolic pathways were detected, with the concentrations of 4 metabolites significantly reduced and 14 metabolites significantly increased in TR26. The concentration of the byproduct L-lactate was sharply reduced due to the deletion of the lactate dehydrogenase gene *ldh*. Additionally, the shikimate concentration in TR26 increased by 117-fold (Fig. S1), which is an intermediate for L-tryptophan synthesis. This suggests requirement to further optimize shikimate pathway, despite the enhanced flux from shikimate to

**Fig. 1.**
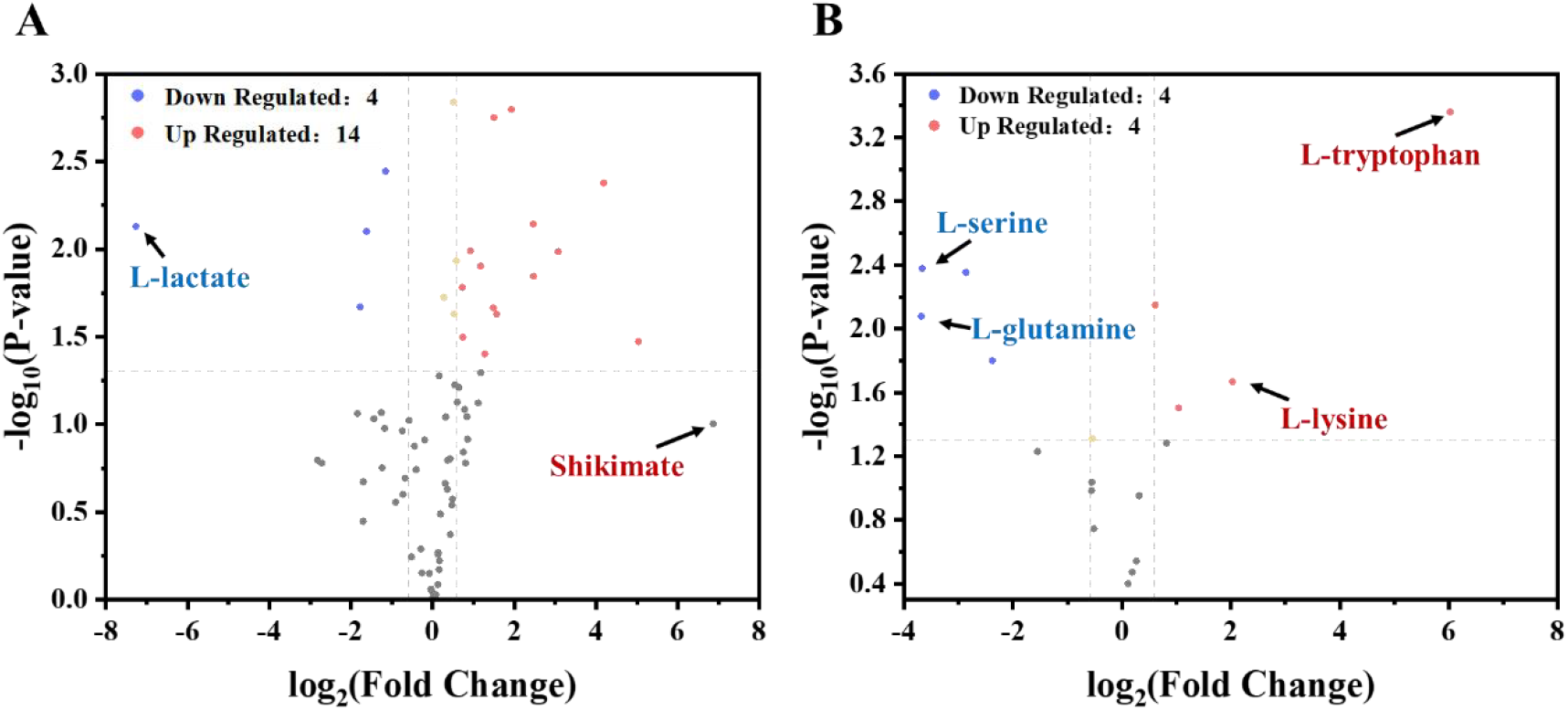
Comparative metabolome analysis of TR26 and MB001. (A) Concentration changes of intracellular metabolites within central metabolic pathways. (B) Concentration changes of intracellular amino acids. Metabolites with average fold change ≥1.5, t-test p ≤0.05 were considered significantly different.

L-tryptophan achieved through the overexpression of *aroC* and *aroD* [17]. On the other hand, intracellular concentrations of L-tryptophan precursors, phosphoenolpyruvate (PEP) and erythrose-4-phosphate (E4P), slightly improved in TR26 (Fig. S1), implying no significant shortage in supply of these precursors. Notably, TR26 exhibited a generalized decline in pyruvate and tricarboxylic acid (TCA) cycle metabolite concentrations (Fig. S1), suggesting a redirection of carbon flux from biomass accumulation toward L-tryptophan synthesis.

Regarding the intracellular concentrations of amino acids, TR26 exhibited significant increase in 4 amino acids and decrease in 4 amino acids compared with MB001 (Fig. 1B). The intracellular concentration of L-tryptophan in TR26 was 65-fold higher than in MB001 (Fig. S1), yet exhibited significant decrease compared with the previous strain TR13. This confirms the effective enhancement of product efflux due to introduction of the L-tryptophan exporter protein [17]. Notably, the concentration of L-lysine increased by 4.1-fold (Fig. 1B), suggesting the necessity for targeted attenuation of its biosynthetic pathway. On the other hand, the accumulation of L-tyrosine and L-phenylalanine in TR26 showed no improvement compared with MB001, which could be attributed to deletion of the *pat* gene that alleviated carbon flux competition among aromatic amino acids. However, significant decline in L-serine and L-glutamine concentrations in TR26, which are precursors for L-tryptophan synthesis, implied the emergence of new bottlenecks for product accumulation.

### 3.2. Transcriptome analysis of *C. glutamicum* TR26 and MB001

To comprehensively analyze the mechanisms underlying L-tryptophan overproduction in the engineered *C. glutamicum* TR26 and provide potential strategies for further strain optimization, comparative transcriptome analysis was performed between strains TR26 and the wild-type MB001. As shown in Fig. 2A, 455 genes were significantly down-regulated and 524 genes were significantly up-regulated in the exponential phase in strain TR26 compared with MB001. In the stationary phase, on the other hand, 392 genes were significantly down-regulated and 343 genes were up-regulated in TR26 (Fig. 2B). KEGG analysis revealed that the DEGs in exponential phase were primarily enriched in pathways related to microbial metabolism in diverse environments, biosynthesis of amino acids, and ABC transporters (Fig. S2). This indicates significant changes in amino acid biosynthesis and extensive influence of the engineering strategies on cell metabolism in TR26. In the stationary phase, KEGG analysis suggested that the DEGs were predominantly enriched in pathways involved in biosynthesis of amino acids, ribosome, and two-component systems (Fig. S2). This implys complex metabolic regulation in TR26 and differences in transcription profiles during distinct growth phases, in addition to general alterations in amino acid metabolism, especially L-tryptophan synthesis, induced by genetic modifications.

**Fig. 2.**
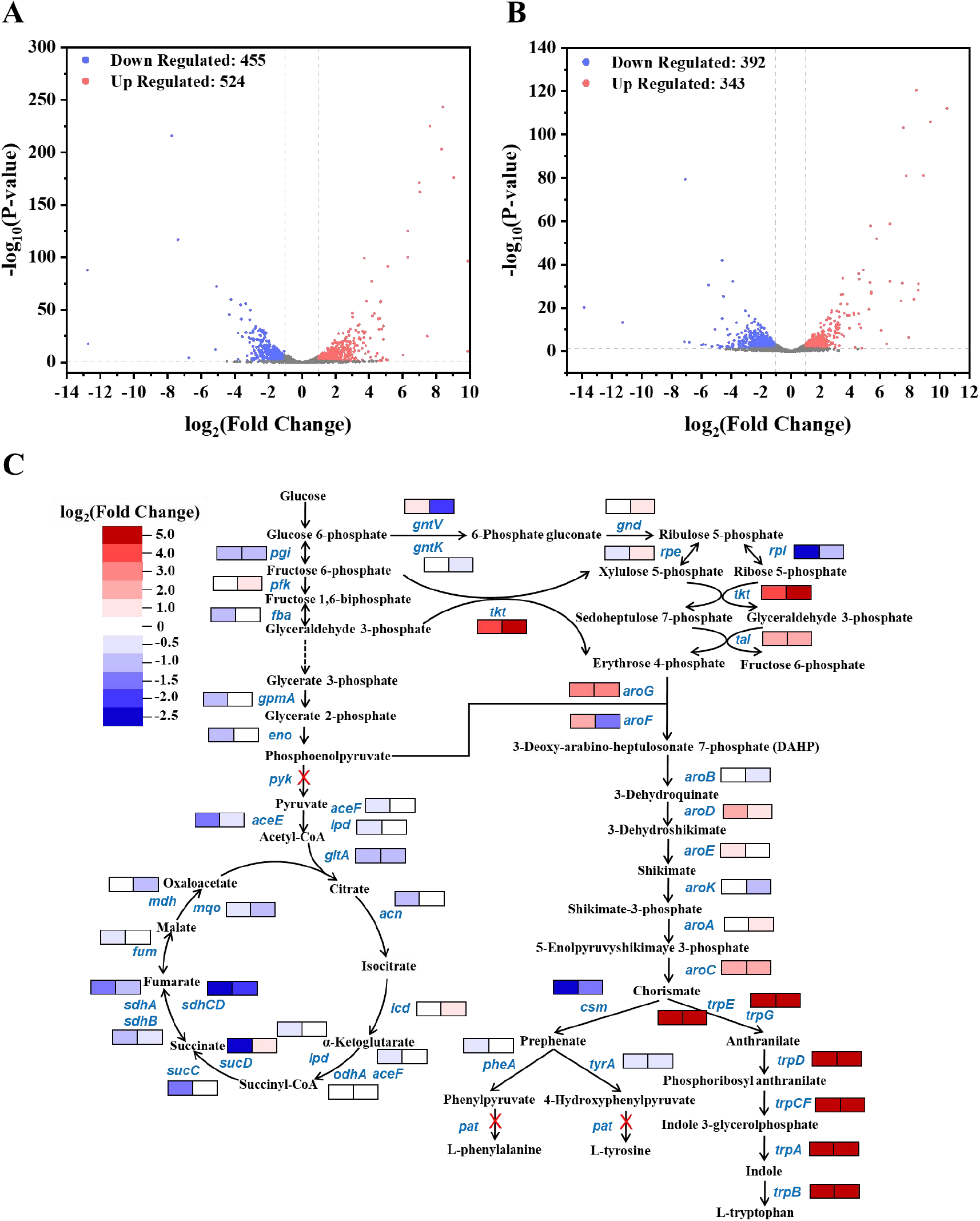
Transcriptome analysis between strains TR26 and MB001. (A) Global gene expression characteristics of strains TR26 and MB001 in the exponential phase. (B) Global gene expression characteristics of strains TR26 and MB001 in the stationary phase. (C) Gene expression profiles in strain TR26 relative to wild-type MB001 within central metabolic pathways and aromatic amino acid synthesis pathway. The right and left boxes represent variations in gene expression levels during the exponential phase and stationary phase, respectively. “X” colored in red indicates genes deleted in TR26. Genes with average fold changes ≥2, t-test p ≤0.05 were considered significantly different.

As shown in Fig. 2C, analysis of genes in central metabolic pathways revealed that most genes in the glycolysis pathway and TCA cycle were down-regulated during the exponential phase in TR26. This indicaties a shift of metabolic fluxes from biomass accumulation to product synthesis, which was also suggested by the significant up-regulation of the *trp* operon genes (*trpEGDCFBA*) during both growth stages. However, in the stationary phase, expression level variations of genes in glycolysis pathway and TCA cycle between TR26 and MB001 were generally alleviated (Fig. 2C), implying the possibility of carbon loss due to enhanced TCA cycle metabolism. Therefore, dynamic regulation of the key genes within central metabolic pathways could be an effective strategy to improve L-tryptophan accumulation at the late stage of fermentation. Notably, the transcription level of *tkt*, which was overexpressed in TR26, was increased by 14.6- and 23.7-fold in the exponential and stationary phase, respectively (Fig. 2C). This might substantially improve E4P supply for L-tryptophan overproduction.

Further analysis of genes within the shikimate and aromatic amino acid synthesis pathways revealed that the expression levels of *aroC* and *aroD* were significantly improved in TR26 during both growth stages. This confirms the crucial roles of these two genes in optimizing the shikimate pathway for L-tryptophan production in *C. glutamicum* [17]. On the other hand, genes in the L-tyrosine and L-phenylalanine synthesis pathways, specifically *csm, pheA*, and *tyrA*, were significantly down-regulated in TR26. This indicates effective repression of the competing pathways, which was also reflected in the metabolome analysis and represents one of the key mechanisms of L-tryptophan overproduction in *C. glutamicum* TR26.

### 3.3. Screening of novel genetic targets for L-tryptophan production

The DEGs between *C. glutamicum* strains TR26 and MB001 could involve genes related to L-tryptophan synthesis, which could be engineered to further enhance L-tryptophan production in TR26. It was shown that 214 genes were down-regulated, and 183 genes were up-regulated in both the exponential and stationary phases between TR26 and MB001. Functional analysis of these genes identified 43 candidates related to L-tryptophan biosynthesis, which were involved in transcriptional regulation, redox reaction, energy metabolism, stress response, amino acid and aromatic compound metabolism, or cell membrane (Table 1).

**Table 1.**
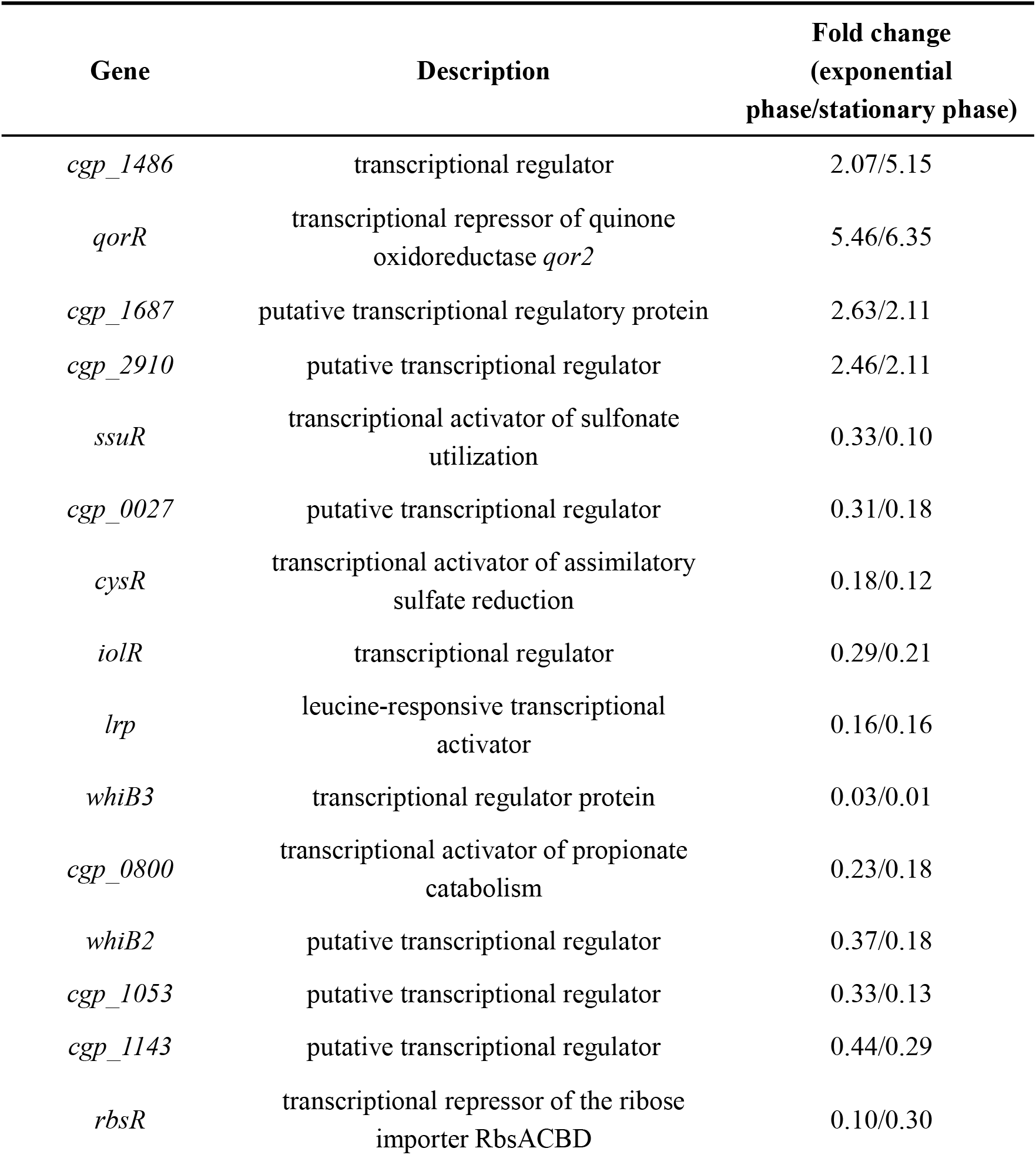
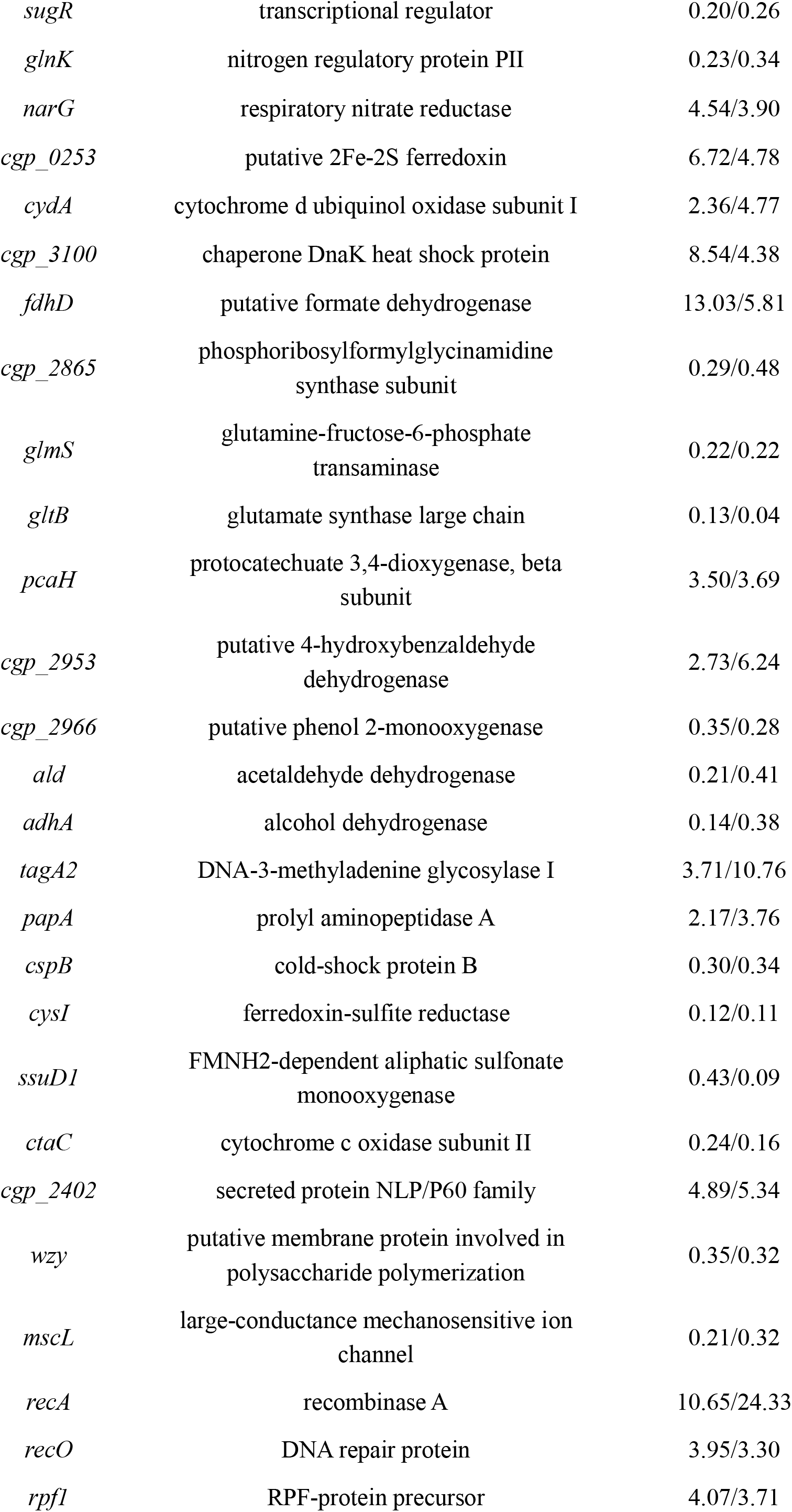
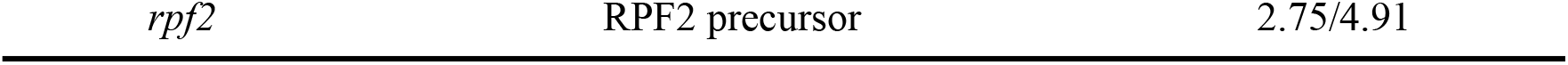
Candidate genetic targets related to L-tryptophan synthesis.

To investigate the effect of down-regulating these candidate genes on L-tryptophan production, we employed the CRISPR interference (CRISPRi) system. Specific sgRNA targeting each candidate gene was designed and ligated to the pDSG plasmid, which contains the dCas9 protein and sgRNA cassette. The plasmids were subsequently introduced into strain TR26, generating an arrayed cell library with repressed candidate genes. Strain TR26 containing the pDSG plasmid with non-targeting sgRNA served as the negative control (NC). As shown in Fig. 3, the repression of *glnK, cydA, papA, ssuD1*, and *cgp_2402* resulted in increased L-tryptophan titers (>8%) compared to the control group. Conversly, strains with down-regulated *qorR, cgp_1687, cysR, lrp, whiB3, rbsR, sugR, narG, cgp_0253, cgp_2865*, and *cysI* exhibited decreased L-tryptophan accumulation (>12%). Additionally, the repression of *cysR, whiB2, cgp_2865*, and *cysI* led to significant reductions in biomass accumulation (Fig. S3), which could negatively influence cell physiology.

**Fig. 3.**
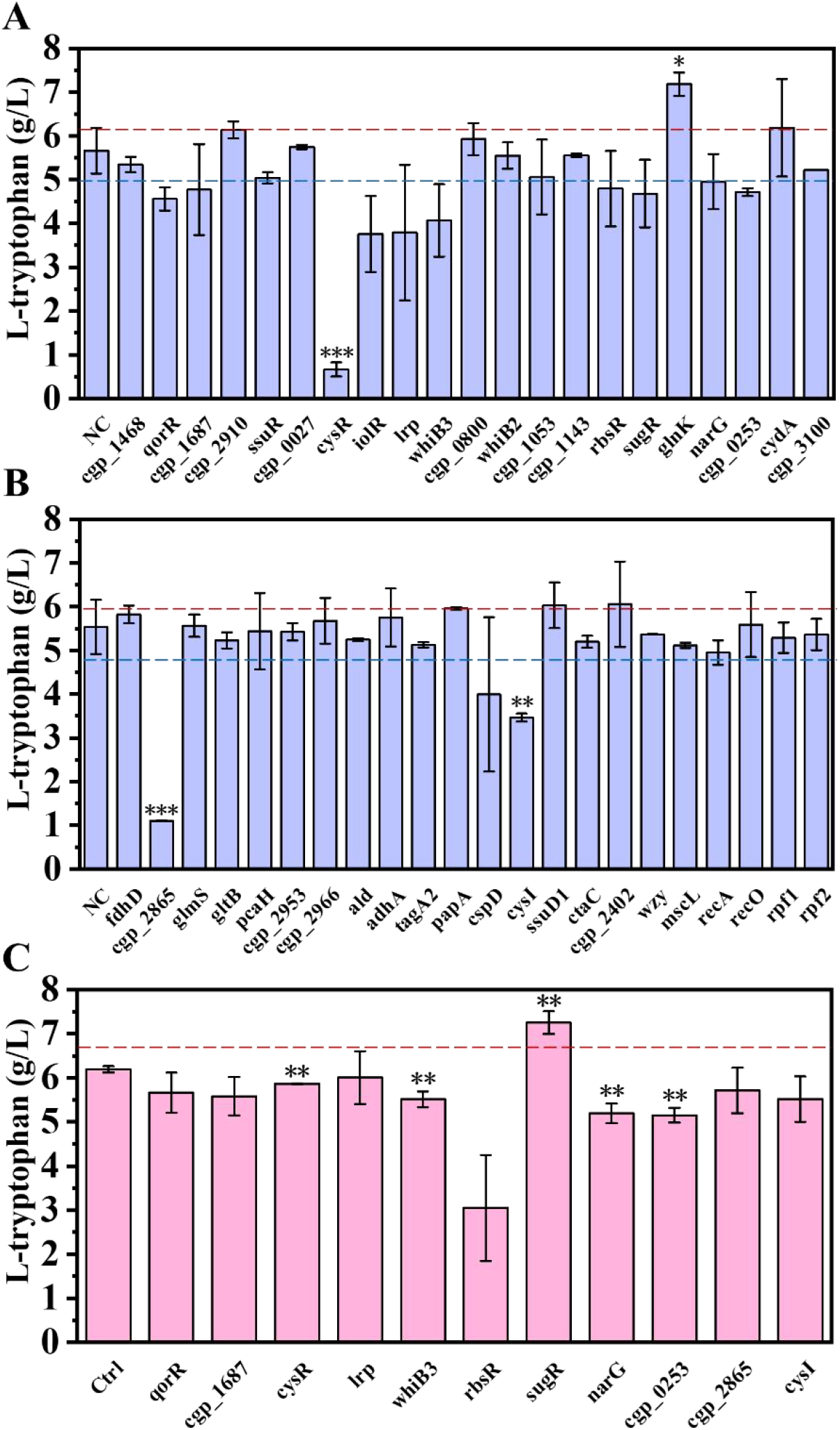
Screening of novel genetic targets for L-tryptophan overproduction. (A)(B) L-tryptophan titers of strains with down-regulated target genes in 48-well microtiter plates. (C) L-tryptophan titers of strains with up-regulated target genes in 48-well microtiter plates. NC, TR26 containing pDSG plasmid with non-targeting sgRNA. Ctrl, TR26 containing empty pEC-K18mob2 plasmid. Two tailed t-tests indicate statistical significance compared to the control group. *P<0.1, **P<0.05.

Furthermore, we utilized the plasmid pEC-K18mob2 to overexpress the candidate genes whose repression resulted in decreased L-tryptophan production. As shown in Fig. 3C, the strain overexpressing *sugR* exhibited significant improvement in L-tryptophan titer, along with a notable increase in biomass accumulation (Fig. S3). In conclusion, down-regulation of genes *glnK, cydA, papA, ssuD1*, and *cgp_2402*, as well as up-regulation of gene *sugR* could be beneficial for enhancing L-tryptophan synthesis in *C. glutamicum*.

### 3.4. Verification of novel genetic targets by shake-flask fermentation

To verify the effects of down- or up-regulating the novel genetic targets in *C. glutamicum*, and to improve L-tryptophan production in strain TR26, shake-flask fermentations were conducted. As shown in Fig. 4A, among the strains with down-regulated target genes, strains with repressed *glnK* and *cydA* exhibited increased L-tryptophan titers. Notably, the L-tryptophan concentration of the *glnK*-repressed strain was consistently higher than that of the control group (NC) throughout the fermentation process, with an ultimate improvement of 6.7% in product titer, indicating significantly enhanced L-tryptophan producing capacity. Additionally, the OD_600_ values of the strains with down-regulated *glnK* and *cydA* were comparable to the control group (Fig. 4B), suggesting that the down-regulation of the two genes posed no significant impair to cell physiology and growth. Furthermore, the repression of *glnK* and *cydA* resulted in a reduction of acetate accumulation by 26.0% and 41.1%, respectively (Fig. S4), implying improved metabolic balance and redirection of metabolic flux from byproduct synthesis to L-tryptophan production.

**Fig. 4.**
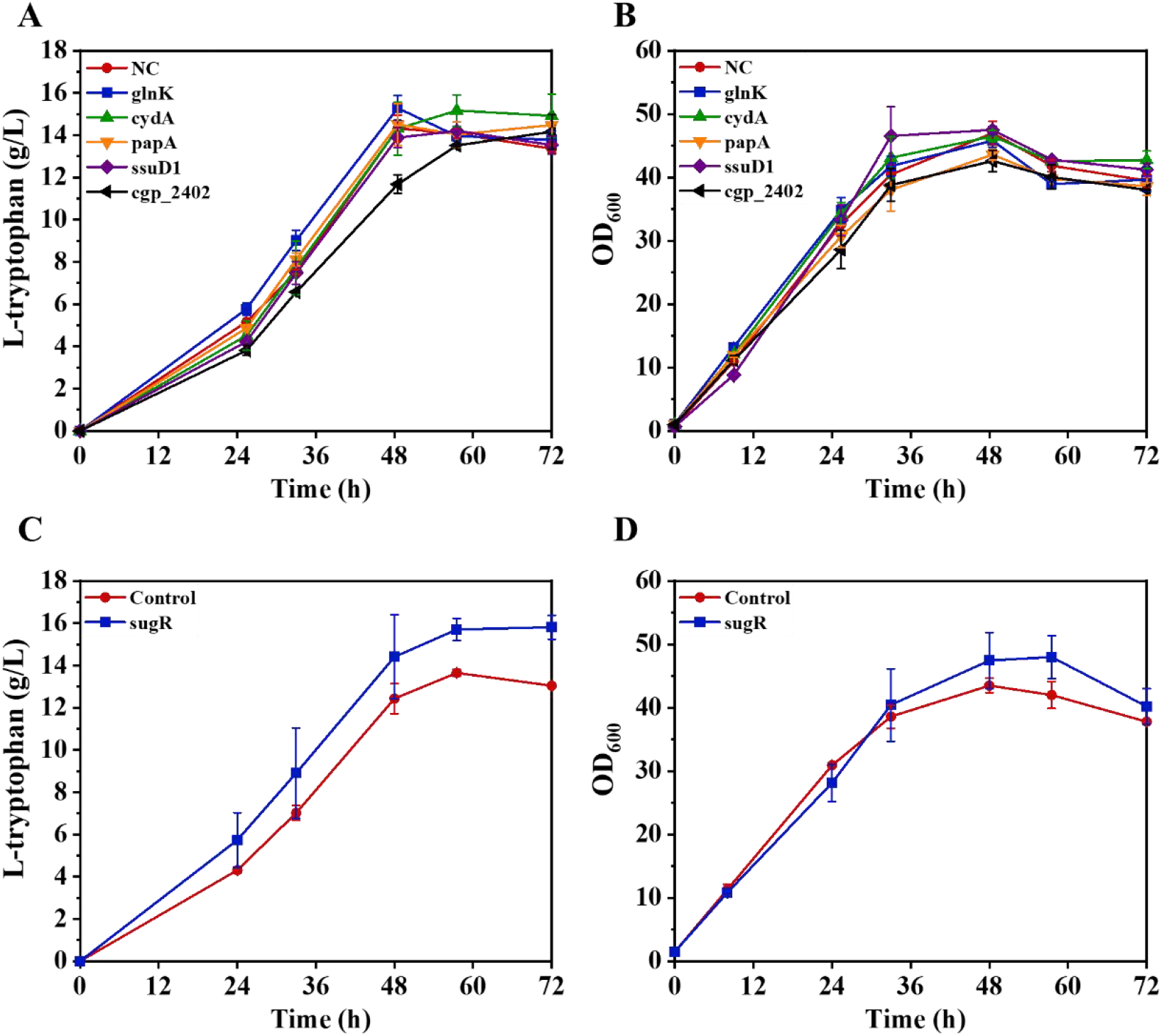
Shake-flask fermentations of the engineered strains. (A) (C) L-tryptophan production. (B) (D) Cell growth. NC, TR26 containing pDSG plasmid with non-targeting sgRNA. Control, TR26 containing empty pEC-K18mob2 plasmid.

On the other hand, the overexpression of *sugR* significantly enhanced L-tryptophan synthesis in strain TR26 in shake-flask fermentation (Fig. 4C). The L-tryptophan titer of the *sugR*-overexpressed strain was improved by 20.9% (Fig. 4C), along with increased biomass accumulation (Fig. 4D) and eliminated acetate secretion (Fig. S4), demonstrating optimized cell metabolism and enhanced L-tryptophan production.

### 3.5. Regulation mechanism of *glnK* on L-tryptophan production

It has been demonstrated that the down-regulation of gene *glnK* is beneficial for L-tryptophan production in strain TR26. However, the regulation mechanism of *glnK* on L-tryptophan synthesis requires further elucidation. GlnK is a crucial component of the nitrogen regulation system in *C. glutamicum*, which functions by interacting with the global transcription factor AmtR. In conditions with limited nitrogen sources, GlnK becomes adenylylated by GlnD and was enabled to bind AmtR and derepress the AmtR regulons, including genes involved in ammonium uptake and assimilation (e.g., *amt, amtB, glnA*, and *gltB*), urea transport and metabolism (e.g., *ureABCDEFG*), and signal transduction (e.g., *glnK* and *glnD*) (Fig. 5A) [49,50]. Comparative analysis of gene transcription profiles between the TR26 strain containing non-targeting sgRNA plasmid (NC, the control group) and the strain with down-regulated *glnK* gene revealed that the expression repression of *glnK* resulted in significant down-regulation of the AmtR regulons (Fig. 5B and C), consistent with previous findings. Notably, the transcription level of the glutamine synthase gene *glnA* was reduced by 51.1% (Fig. 5C). Therefore, the down-regulation of *glnK* in strain TR26 could lead to decreased nitrogen uptake and assimilation efficiency, as well as reduced L-glutamine synthesis capacity.

**Fig. 5.**
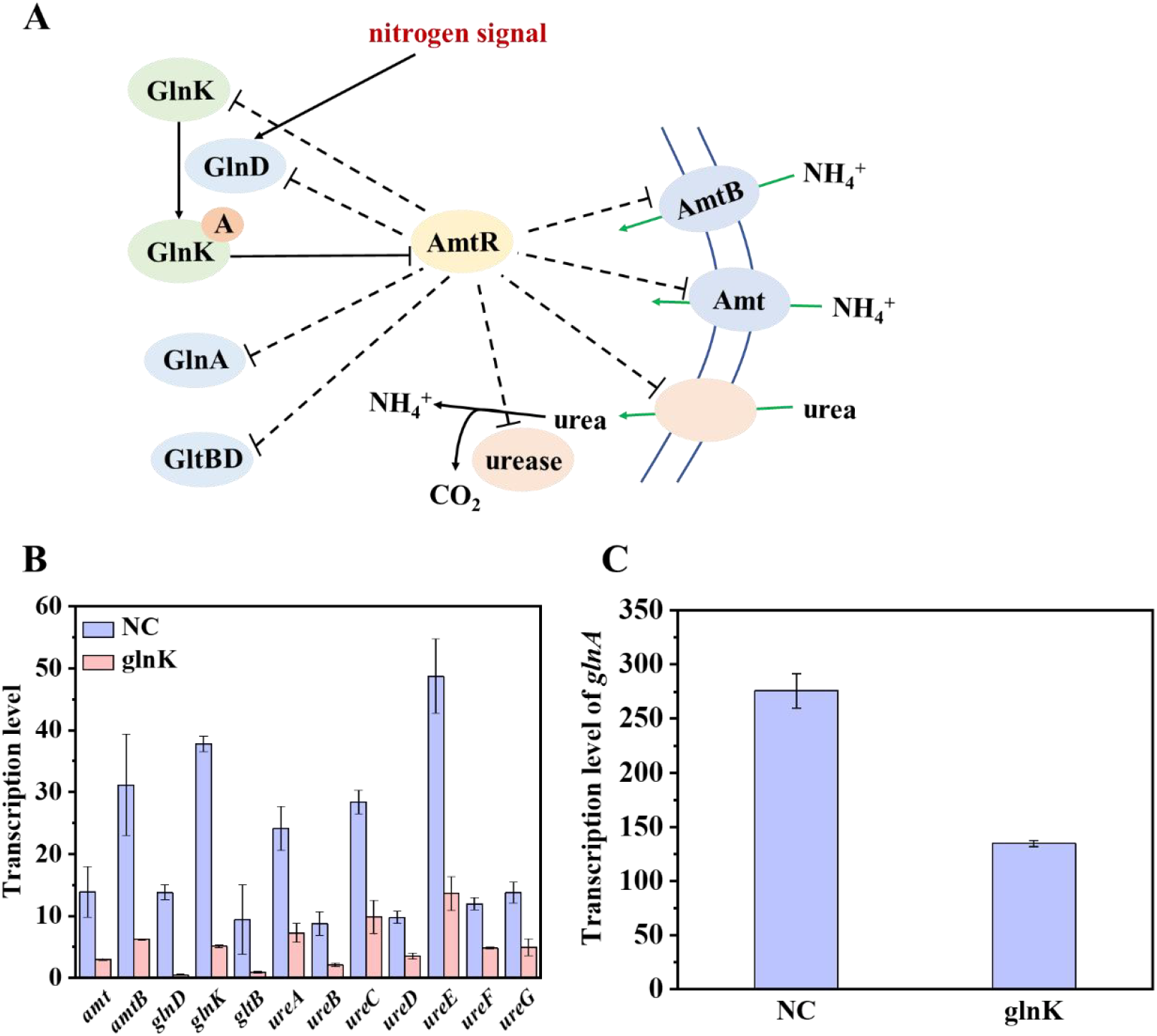
Regulation mechanism of GlnK. (A) The nitrogen regulation network mediated by GlnK in *C. glutamicum*. Arrowheads indicate signal transduction; solid lines with blunt ends indicate regulation of enzyme activity; dashed lines indicate expression regulation. (B) (C) Transcription levels of genes related to nitrogen utilization. NC, TR26 containing pDSG plasmid with non-targeting sgRNA; glnK, TR26 with down-regulated *glnK* gene mediated by CRISPRi.

To further investigate the effects of *glnK* down-regulation on cell metabolism, intracellular amino acid concentrations were analyzed. As shown in Table 2, in strain TR26 with repressed *glnK*, the concentrations of L-glutamate and L-glutamine were reduced to 87% and 34% of the control group, respectively. In addition, L-arginine, L-proline, and L-tyrosine, which are nitrogen-rich amino acids, also exhibited decreased concentrations (Table 2). This suggested that the down-regulation of *glnK* might enhance L-tryptophan production by reducing metabolic fluxes of the competing pathways. On the other hand, considering the fermentation medium contains abundant nitrogen sources (i.e., ammonium and urea), which can be transported into the cell through diffusion, the down-regulation of nitrogen uptake-related genes (Fig. 5B) may not limit intracellular nitrogen availability. Instead, it could allocate more resources to metabolic pathways supporting L-tryptophan synthesis rather than nitrogen uptake, thereby promoting L-tryptophan production.

**Table 2.** The effect of *glnK* repression on intracellular amino acid concentrations.

| Amino acid | Fold change | t-test P |
| --- | --- | --- |
| L-glutamate | 0.87 | 0.54 |
| L-glutamine | 0.34 | 0.01 |
| L-arginine | 0.70 | 0.001 |
| L-proline | 0.50 | 0.001 |
| L-tyrosine | 0.41 | 0.06 |

### 3.6. Regulation mechanism of *sugR* on L-tryptophan production

The novel genetic target for L-tryptophan production identified in this research, *sugR*, is a global transcriptional regulator involved in sugar uptake and metabolism in *C. glutamicum*. SugR, encoded by the *sugR* gene, functions as transcriptional repressor of the phosphotransferase system (PTS) encoding genes including *ptsI, ptsH, ptsF, ptsG*, and *ptsS* [51], as well as glycolysis pathway genes such as *pfk, fba*, and *eno* [52] (Fig. 6A). Comparative transcriptome analysis demonstrated that the overexpression of *sugR* significantly reduced the transcription levels of these genes, as expected (Fig. 6B and C). Conversly, the non-PTS genes *iolT1* and *glk* were up-regulated by 3.7- and 1.6-fold, respectively, in the *sugR*-overexpressed strain, suggesting activation of the non-PTS to compensate for sugar uptake.

**Fig. 6.**
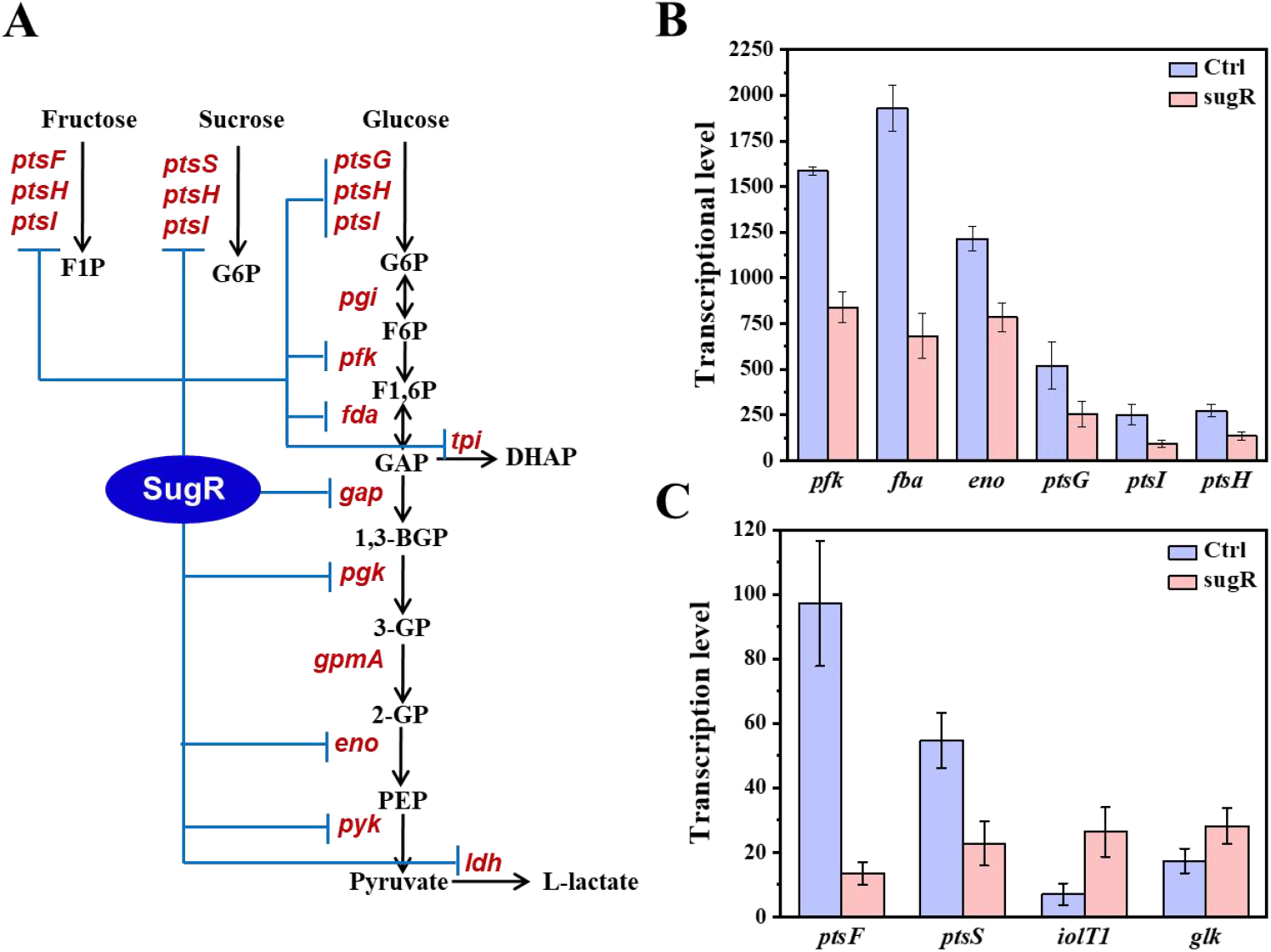
Regulation mechanism of SugR. (A) Genes regulated by SugR in *C. glutamicum*. (B) (C) Transcription level variations of genes related to *sugR* regulation. Ctrl, TR26 containing empty pEC-K18mob2 plasmid.; sugR, TR26 with up-regulated *sugR* gene.

In *C. glutamicum*, the PTS is the primary sugar uptake system which requires PEP for phosphorylation, thus reducing the PEP availability for L-tryptophan synthesis. In contrast, the non-PTS depends on ATP instead of PEP for phosphorylation and is typically inactive. Therefore, by repressing PTS and activating non-PTS, the overexpression of *sugR* could reduce PEP consumption in sugar uptake processes, thereby effectively improving the supply of precursor PEP and enhancing the production of L-tryptophan.

## 4. Conclusion

In conclusion, this study systematically investigated the mechanisms and potential bottlenecks of L-tryptophan overproduction in *C. glutamicum* TR26, and identified novel genetic targets related to L-tryptophan synthesis. By integrating comparative metabolome and transcriptome analyses between strains TR26 and MB001, it was revealed that several key factors contributed to L-tryptophan overproduction in *C. glutamicum*, including the redirection of metabolic fluxes from TCA cycle to L-tryptophan synthesis, enhanced L-tryptophan export, optimized PEP and E4P supply, debottlenecked shikimate pathway, as well as repressed synthesis of L-tyrosine and L-phenylalanine. However, potential bottlenecks such as carbon loss during the late fermentation stage and limited supply of L-serine and L-glutamine were idintified in strain TR26. Additionally, by systematically down- and up-regulating key differentially expressed genes, two novel genetic targets, *glnK* and *sugR*, were identified. The repression of *glnK* in strain TR26, which could affect the uptake and assimilation of nitrogen sources, as well as optimize cell resource allocation for L-tryptophan synthesis, resulted in a 6.7% increase in L-tryptophan titer and a 26.0% reduction in acetate accumulation. Meanwhile, the overexpression of *sugR*, which could enhance PEP supply for L-tryptophan synthesis by reducing PEP consumption in sugar transport processes, led to a 20.9% improvement in L-tryptophan titer and eliminated acetate accumulation in TR26. Future efforts to enhance L-tryptophan production in strain TR26 can be focused on genome engineering of the novel targets, as well as dynamic regulation of central metabolic pathways to mitigate carbon loss from the TCA cycle in the late stages of fermentation.

## Acknowledgments

This work was supported by the National Key R&D Program of China (No. 2021YFC2100900), the National Natural Science Foundation of China (Grant Nos. 21938004, 22078172), and Tsinghua University Initiative Scientific Research Program (No. 20223080016).

## Author Contributions

Yufei Dong: Methodology, Investigation, Data curation, Validation, Writing-original draft preparation. Rongsheng Gao: Formal analysis, Data curation. Nan Qin: Methodology, Investigation. Kunyu Liu: Methodology, Investigation. Youmeng Liu: Methodology, Investigation. Zhen Chen: Conceptualization, Investigation, Writing-review & editing, Funding acquisition, Resources, Supervision, Project administration.

