## Supplementary Information for "Transcriptome and metabolome analyses reveal novel genetic targets for L-tryptophan overproduction in *Corynebacterium glutamicum*"

**Table S1. The sgRNA sequences of the candidate genes**

| Gene | SgRNA sequence (5’-3’) |
|---|---|
| *cgp_1486* | TAAAACCTTAATACCGCTCT |
| *qorR* | ATTTGCCGGTTGCTGAATAG |
| *cgp_1687* | CACCTGGGTCAAAAACTTCC |
| *cgp_2910* | CCTTGAGATACCTAGTTTCG |
| *ssuR* | TACGCGCGCGACGGGTAGGT |
| *cgp_0027* | CTTCAAGCGGTCCAAGCGTG |
| *cysR* | GCAGAGAAGTGGGCCTGATT |
| *iolR* | GGCTGGCCAAATGGGAGCTT |
| *lrp* | ACCCTCCTCAAGCAAGGTCC |
| *whiB3* | CAGAAGTCTGCATTTGGCCC |
| *cgp_0800* | ATTGGGTGAGCTGATGGGTT |
| *whiB2* | ATCCAAAGTCATCTCCAGTG |
| *cgp_1053* | CCGCAACCGTGAACGCCTCA |
| *cgp_1143* | GATTCGAGCCGCTAAATCAA |
| *rbsR* | ACGGTAGTTCAGCTTTTGAG |
| *sugR* | TTAATGAGGCAATCTGTCGA |
| *glnK* | GGTAAACGGCTTGACAATTG |
| *narG* | GTAGTCATGGATTGGCCTTT |
| *cgp_0253* | GGGTAGGTCGGCCTGGAGGA |
| *cydA* | GCGCTAAGCCAATGGTCAGT |
| *cgp_3100* | TGGCTCGCCGCCTTCAAGTA |
| *fdhD* | CGTCAGTGGACACAACGCGT |
| *cgp_2865* | ATTGACAACTACACGGGCCA |
| *glmS* | CATCTAGAGCAAAGTAATCA |
| *gltB* | GAGTCCTTGTGGTTTCATGC |
| *pcaH* | GGAGTATTCGCCTCCCGTCG |
| *cgp_2953* | TTGTTTGCTTTCGGCGATTA |
| *cgp_2966* | GCGCATCTCCATTGGGTCAC |
| *ald* | ATAGTTAACGATCGAGCCTT |

| | |
|---|---|
| *adhA* | TGGAAGGTCAATATCCTTCA |
| *tagA2* | GCGAACCATAATTTGAGCAT |
| *papA* | TGCTTCCCAGGTTACGCGCA |
| *cspB* | TGAAGCCGAAGCCCTTCTCT |
| *cysI* | TTCGGGCTTAGGCTTCCTGG |
| *ssuD1* | AGATTCAAAACCGTTGGTCT |
| *ctaC* | TCCAAGCACACCGCCAGGAG |
| *cgp_2402* | TTGCGTTTGAATTGTTGCGA |
| *wzy* | AGCTAAAGGCTCACGAAGCA |
| *mscL* | GGAGAATGCTGTCACGATAG |
| *recA* | CTGACGATCATTCCCCTTGG |
| *recO* | AAGTTTTGACTACTAGCGCG |
| *rpf1* | CTGTGCGAGGCGATCCCAGT |
| *rpf2* | GTTGATCCGTGACTTCTGAT |

**Table S2. Primers used for target gene amplification**

| Overexpressed gene | Primer name | Sequence (5'-3') |
|---|---|---|
| *qorR* | H1-F | accatgattacgaaaggaggttgtcatggatatttcccctattcagcaaccg |
| | H1-R | ttctctcatccgccaaaacagccacttagtgctgcttatcgtagatttcttgcg |
| *cgp_1687* | H2-F | accatgattacgaaaggaggttgtcatgatgtcggatcggaaggaagatctc |
| | H2-R | ttctctcatccgccaaaacagccacttacaccacctcgttttcgactttgc |
| *cysR* | H3-F | accatgattacgaaaggaggttgtcatgcccaatcaggcccacttc |
| | H3-R | ttctctcatccgccaaaacagccacttagggtacgagagtaagtggctcg |
| *lrp* | H4-F | accatgattacgaaaggaggttgtcatgaagctagattccattgatcgcgc |
| | H4-R | ttctctcatccgccaaaacagccactcacacctgggggcgagctg |
| *whiB3* | H5-F | accatgattacgaaaggaggttgtcatgacattgcctcaccagcttcc |
| | H5-R | ttctctcatccgccaaaacagccacttaaactgctactggttgcttgcgc |
| *rbsR* | H6-F | accatgattacgaaaggaggttgtcatggcttccgaaacctccagc |
| | H6-R | ttctctcatccgccaaaacagccactcagagttcaccatctaggctcacc |
| *sugR* | H7-F | accatgattacgaaaggaggttgtcatgtacgcagaggagcgcc |
| | H7-R | ttctctcatccgccaaaacagccactcattctgcaatcacaacttctacatcgc |
| *narG* | H8-F | accatgattacgaaaggaggttgtcatgactacaactacttcttctgggaagtcttc |
| | H8-R | ttctctcatccgccaaaacagccacttaggcgccgtgccatttcg |
| *cgp_0253* | H9-F | accatgattacgaaaggaggttgtcatggccctgcacacgcac |
| | H9-R | ttctctcatccgccaaaacagccacctagtcatcgaaggaacagtggtatggc |
| *cgp_2865* | H10-F | accatgattacgaaaggaggttgtcgtggcccgtgtagttgtcaatgtc |
| | H10-R | ttctctcatccgccaaaacagccacttatgcaactactgcgttagctccaac |
| *cysI* | H11-F | accatgattacgaaaggaggttgtcatgacaacaaccaccggaagtgc |
| | H11-R | ttctctcatccgccaaaacagccactcatgagtgaagtccgcattctgtc |

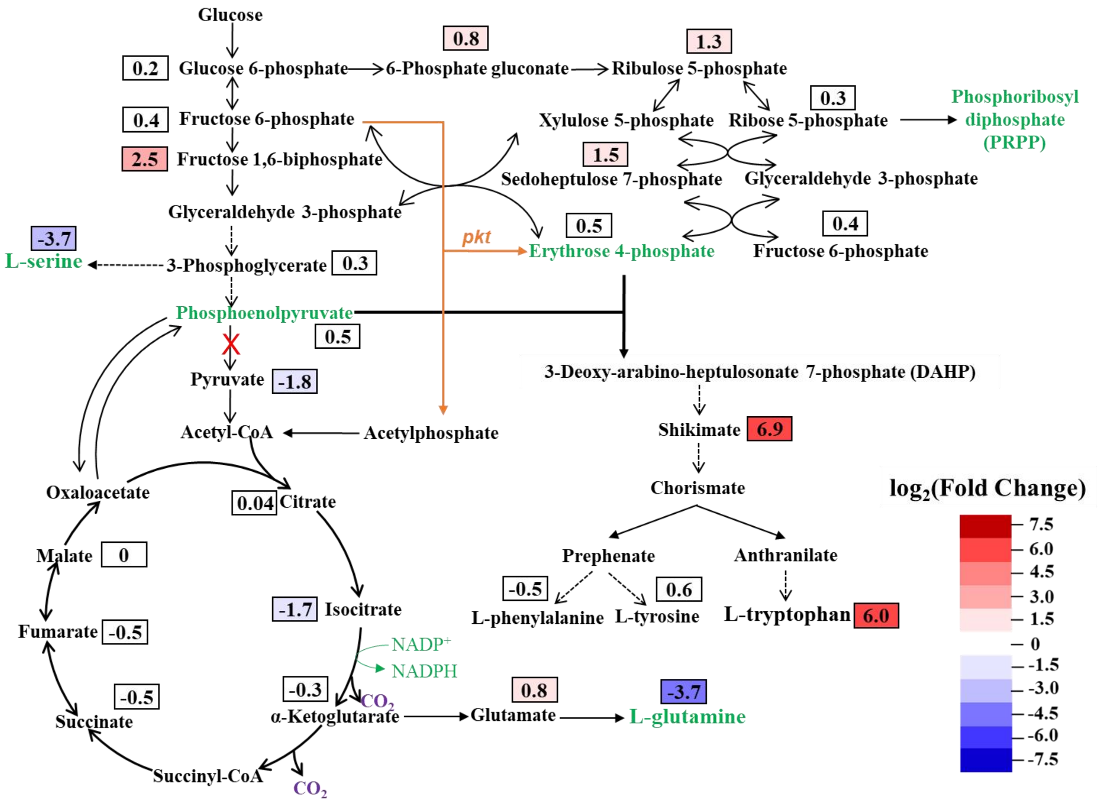


**Fig. S1. Comparative metabolome analysis between *C. glutamicum* strains TR26 and MB001.** The numbers indicate the $log_2$(Fold Change) of the intracellular concentrations of the metabolites in TR26 compared with MB001.

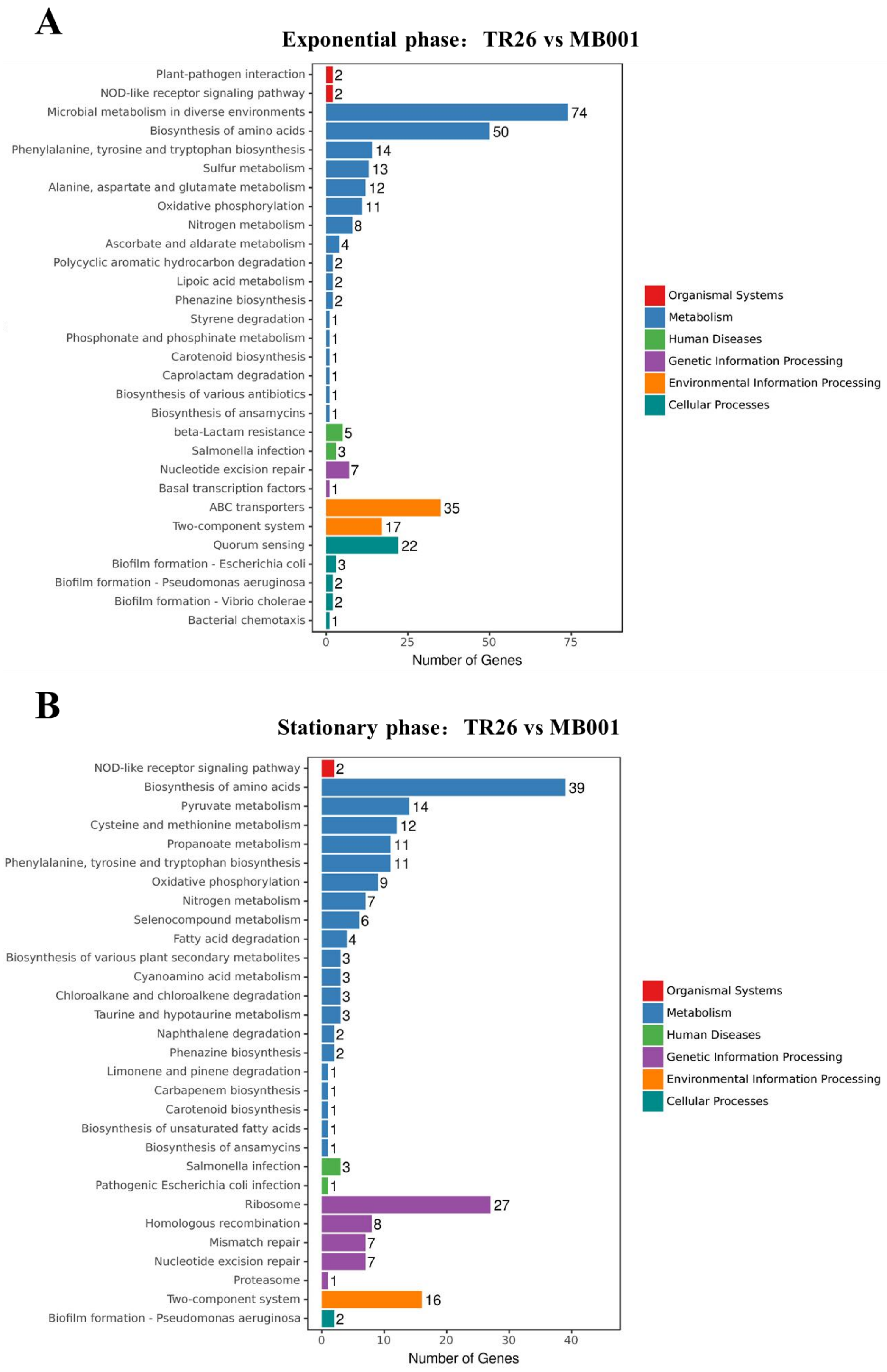


**Fig. S2. KEGG pathway enrichment analysis of differentially expressed genes between *C. glutamicum* strains TR26 and MB001.** (A) Exponential phase. (B) Stationary phase.

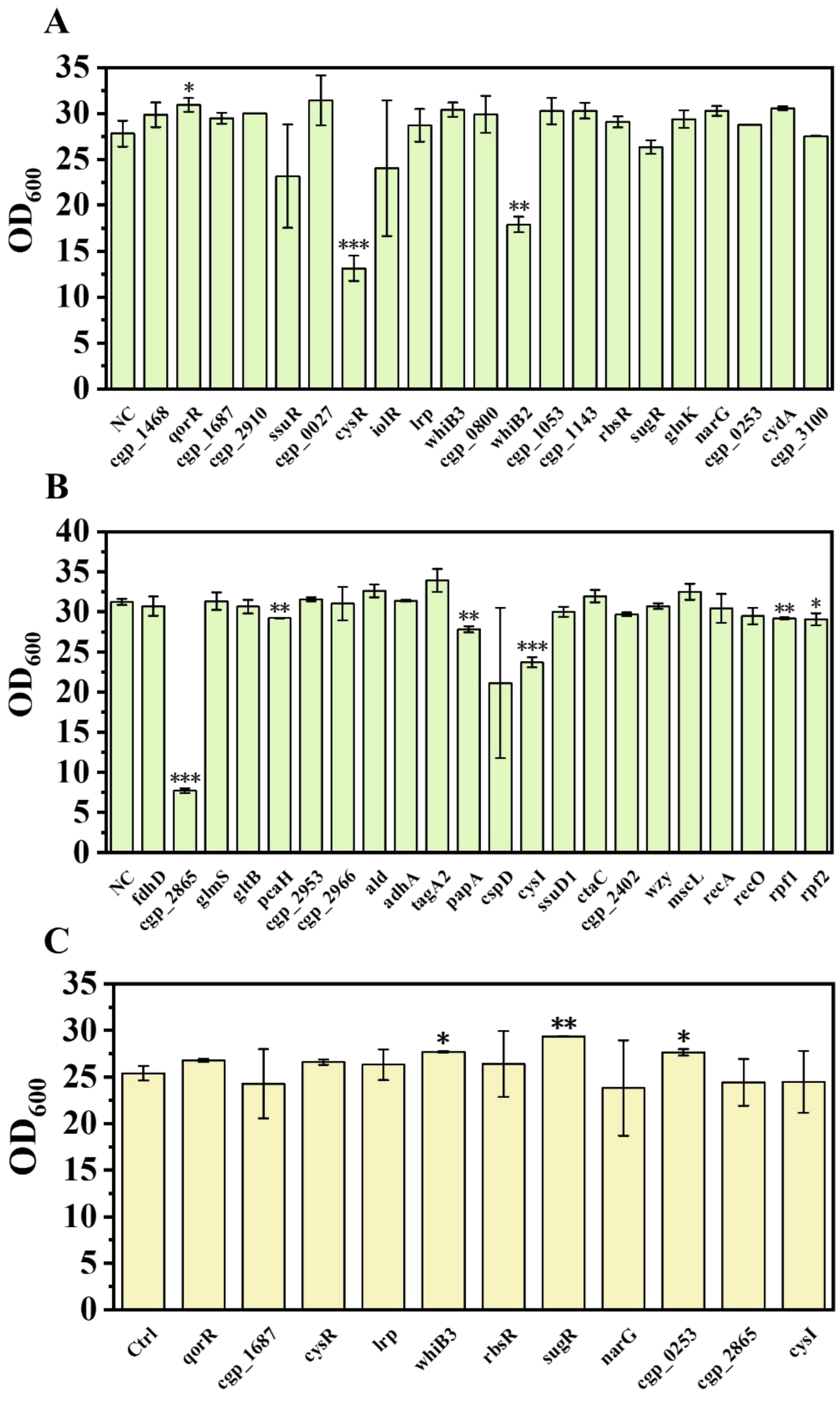


**Fig. S3. Screening of novel genetic targets for L-tryptophan overproduction.** (A) (B) The final $OD_{600}$ of strains with down-regulated target genes in 48-well microtiter plates. (C) The final $OD_{600}$ of strains with up-regulated target genes in 48-well microtiter plates. NC, TR26 containing pDSG plasmid with non-targeting sgRNA. Ctrl, TR26 containing empty pEC-K18mob2 plasmid. Two tailed t-tests indicate statistical significance compared to the control group. *P<0.1, **P<0.05, ***P<0.01.

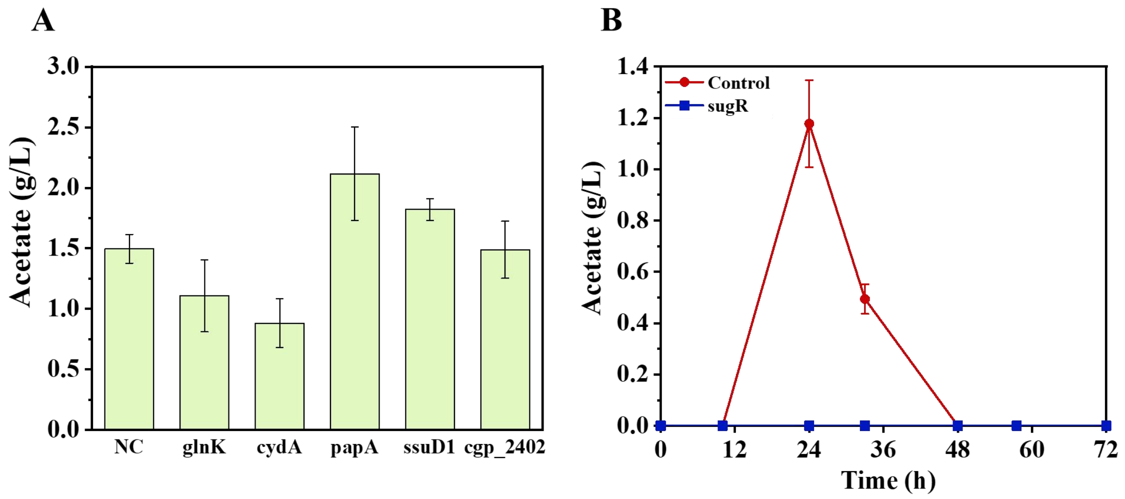


**Fig. S4. Acetate accumulations of the engineered strains during shake-flask fermentations.** (A) Strains with down-regulated genetic targets. (B) Strains with up-regulated genetic target. NC, TR26 containing pDSG plasmid with non-targeting sgRNA. Control, TR26 containing empty pEC-K18mob2 plasmid.
